# Anatomical and Neurochemical Profiles of GABAergic Projection Neurons in the Mouse Inferior Colliculus

**DOI:** 10.1101/2025.09.02.673871

**Authors:** Ryohei Tomioka, Kenta Kobayashi, Wen-Jie Song

## Abstract

The inferior colliculus (IC) is a critical hub for the integration of auditory signals in the midbrain. Although both glutamatergic and GABAergic neurons in the IC project to the medial geniculate body (MGB), the detailed neurocircuitry and neurochemical properties of GABAergic projection neurons remain poorly understood. In this study, glutamate decarboxylase 67 (GAD67)-Cre mice and the viral vector were utilized to selectively visualize the axonal projections of GABAergic neurons in the IC. These neurons project predominantly to the medial division of MGB (MGv) and contralateral IC and minorly to the ventral nucleus of the trapezoid body. Importantly, GABAergic projections from both lemniscal and non-lemniscal regions of the IC primarily targeted the lemniscal division of the MGB, whereas those projections from the external cortex of the IC—part of the non-lemniscal pathway—additionally extended into non- lemniscal MGB subregions. Using a retrograde tracer, a substantial proportion of GABAergic projection neurons targeting the MGB and contralateral IC were positive for several neurochemical markers, implying that some GABAergic neurons send axon collaterals to both targets. Notably, GABAergic neurons constituted 10% of the total neuronal population in the IC, whereas GABAergic neurons accounted for approximately 20% of IC neurons projecting to the MGB or contralateral IC. Our results suggest that GABAergic projection neurons are more involved in the IC–MGB pathway than would be expected based on their proportion in the IC and may exert widespread influence on auditory processing via direct inhibition of the MGv and indirect modulation through suppression of the contralateral IC.

## Introduction

Auditory information ascends to the auditory cortex via a series of brainstem and subcortical nuclei, including the cochlear nucleus (CN), superior olivary complex (SOC), IC, and MGB (Roger and Arnault 1989; Romanski and LeDoux 1993; Kelly et al. 1998; King et al. 2019). The ascending auditory pathway is comprised of two major streams: the lemniscal pathway, which is responsible for transmitting precise auditory information, and the non-lemniscal pathway, which processes both auditory and multimodal information (Malmierca et al. 2015; Liu et al. 2022).

The IC serves as a critical hub, integrating frequency and temporal information from the CN and spatial information from the SOC (Vonderschen and Wagner 2014). Within the IC, the central nucleus (CNIC) is associated with the lemniscal pathway, whereas the external (ECIC) and dorsal (DCIC) cortices are associated with non-lemniscal pathways. These IC subregions contain both glutamatergic and GABAergic neurons, and send parallel excitatory and inhibitory projections to the MGB (Winer et al. 1996; Ito et al. 2009; Mellott et al. 2014a; Beebe et al. 2018; Silveira et al. 2020). This parallel organization of excitatory and inhibitory projections is a unique feature of the auditory pathway, distinguishing it from other ascending sensory pathways, such as the visual system.

The MGB also contains subregions corresponding to lemniscal and non-lemniscal pathways (Ledoux et al., 1987; Anderson and Linden 2011). The MGv is a lemniscal region that primarily receives input from the CNIC, whereas the other MGB subregions comprise non-lemniscal regions and are thought to receive input from the DCIC and ECIC. Although this connectivity is primarily mediated by glutamatergic projection neurons, GABAergic projection neurons are also known to contribute to the IC-MGB connections (Winer et al. 1996; Ito et al. 2009; Mellott et al. 2014a; Beebe et al. 2018; Silveira et al. 2020). However, to the best of our knowledge, the precise connectivity of GABAergic projection neurons in the IC remains unclear, particularly with respect to the specific connections between individual IC and MGB subregions. Additionally, although candidate neurochemical markers for IC GABAergic projection neurons have been reported (Beebe et al. 2018; Silveira et al. 2020), the proportion of total GABAergic projection neurons expressing these markers remains unclear.

In this study, we employed a variety of anatomical techniques to characterize GABAergic projection neurons in the IC. Our findings sought to clarify the inhibitory connections between the IC and MGB subregions, but also characterize their connectivity with other brain regions and their neurochemical profiles.

## Materials and Methods

All surgical procedures were performed in accordance with the National Institutes of Health (NIH) “Guidelines for the Care and Use of Laboratory Animals” (NIH publication 86-23), and all protocols were approved by the Kumamoto University Animal Experimentation Committee. Every effort was made to minimize animal suffering and to reduce the number of animals used.

Adult animals were housed in a ventilated room under standard conditions, including an air-conditioned environment at 20–25°C and 40–60% humidity, with a 12- hour light/dark cycle (23–30 g at the time of injection). GAD67-Cre knock-in mice (n = 16) (Tomioka et al., 2015) and GAD67-green fluorescent protein (GFP) knock-in mice (n = 14) (Tamamaki et al., 2003) were used in experiments.

### Surgical procedures and fixation

Each animal was anesthetized via intraperitoneal injection of a mixture of ketamine (80 mg/kg; Fujita, Tokyo, Japan) and xylazine (8 mg/kg; Elanco, Tokyo, Japan) and placed in a stereotaxic frame (Narishige, Tokyo, Japan). A craniotomy was performed at the relevant skull location, detailed below. All injections were performed under air pressure using a custom-made syringe pump connected to a glass capillary needle (Calibrated micropipettes 2-000-001-90, Drummond Scientific Company, Broomall, PA, USA). To visualize axonal fibers from the IC, 0.2 µl of adeno-associated virus serotype 2 (AAV2)-hSyn1(S)-Flex-tdTomato-T2A-synaptophysin-fused enhanced GFP (sypEGFP) (Oh et al., 2014) (RRID:Addgene_51509; 7.8 × 10^11^ vg/mL purified at the National Institute for Physiological Sciences, Section of Viral Vector Development) was injected into the IC at the following coordinates: CNIC (bregma, −5.02 mm; lateral, 0.5 mm; depth, 1.0 mm from the surface at a 60-degree angle from the horizontal plane in the medio-lateral direction); ECIC (bregma, −5.02 mm; lateral, 1.2 mm; depth, 0.7 mm from the surface at a 60-degree angle from the horizontal plane in the medio-lateral direction); and DCIC (bregma, −5.02 mm; lateral, 0.2 mm; depth, 0.3 mm at a 60- degree angle from the horizontal plane in the medio-lateral direction). The survival time was three weeks. To retrogradely label projection neurons in the IC, we injected 0.02 µl of Fluoro-Gold (FG; 4% dissolved in water; Fluorochrome) into GAD67-GFP mice. The FG was injected into the MGB (bregma, −3.0 mm; lateral, 2.0 mm; depth, 3.0 mm), CNIC (bregma, −5.02 mm; lateral, 0.5 mm; depth, 1.0 mm from the surface at a 60- degree angle from the horizontal plane), or SOC (Interaural, 0.0 mm; lateral, 1.2 mm; depth, 6.0 mm at a 11-degree angle from the horizontal plane in the rostral-caudal direction). The survival time was 2–3 days.

Mice were perfused under deep anesthesia (ketamine, 120 mg/kg; xylazine, 12 mg/kg) with 0.05 M phosphate-buffered saline (PBS; 0.9% [w/v] saline buffered with 0.05 M sodium phosphate, pH 7.4), followed by 4% formaldehyde in PBS. The brains were cryoprotected with 30% sucrose in PBS, and 50 μm thick coronal sections were cut using a cryostat (CM1950, Leica Biosystems, Nussloch, Germany). Histological analyses were conducted according to the mouse brain atlas to identify specific brain structures (Franklin and Paxinos 2007).

### Fluorescent immunostaining

To identify the anatomical or neurochemical profiles, we performed immunostaining using the following primary antibodies: rabbit anti-GFP antibody (Tomioka et al. 2015), mouse anti-Kv3.1b antibody (Developmental Studies Hybridoma Bank, Iowa, IA), rabbit anti-NeuN antibody (abcam, Cambridge, UK), rabbit anti- neuropeptide Y (NPY) antibody (Merck, Darmstadt, Germany), mouse anti- parvalbumin (PV) antibody (Swant, Marly, Switzerland), mouse anti-protein kinase C δ (PKCδ) antibody (BD Transduction Laboratories, Franklin Lakes, NJ). The detailed information for primary antibodies is summarized in Table 1. After blocking with PBS containing 1% normal donkey serum and 0.3% Triton X-100 (PBST) for 1 h at room temperature, the sections were incubated with primary antibodies overnight in PBST at room temperature. The sections were then incubated with respective secondary antibodies labeled with fluorescent dyes at different wavelengths for 2 h at room temperature (Table 2). Images were obtained using a confocal laser-scanning microscope (FV3000; Olympus, Tokyo, Japan) with an appropriate filters for FG (laser wavelength: 405 nm; detection wavelength: 550-650 nm), Alexa 488 (laser wavelength: 488 nm; detection wavelength: 500–540 nm), Alexa 555 (laser wavelength: 561 nm; detection wavelength: 570–620 nm), or Alexa 647 (laser wavelength: 640 nm; detection wavelength: 650–750 nm). Images were merged using Canvas 12 (Canvas GFX; Boston, MA, USA), and some images were displayed in pseudocolor format.

### Image data analysis

Image analysis was performed using a previously described method, with slight modifications (Tomioka et al. 2024). Similar coronal sections were analyzed for the quantitative measurement of synaptophysin-GFP-labeled axonal fibers from each IC subregion. Images were obtained using a confocal laser-scanning microscope (FV3000, Olympus, Tokyo, Japan) with a 40× objective lens at 0.311 µm/pixel. All images were analyzed using MATLAB software (R2019b; MathWorks, MA, USA). Briefly, the boundaries of the MGB subdivisions were defined according to PKCδ immunoreactivity (Tomioka et al. 2023). The area of each subdivision was calculated. The raw images of GFP-positive signals were converted into binary images using one threshold per section. The summed area of GFP-positive signals in each subdivision was calculated. Another parameter the occupancy of GFP-positive signals was computed as the ratio of the summed area of GFP-positive signals to the total area of each MGB subdivision. Friedman’s test and Bonferroni’s post hoc test were used for multiple comparisons. Statistical significance was set at p < 0.05.

## Results

### Axon terminals of GABAergic projection neurons in the IC subregions

Although retrograde tracer studies have shown that GABAergic neurons in the IC project to the MGB (Winer et al. 1996; Ito et al. 2009; Mellott et al. 2014b; Beebe et al. 2018; Silveira et al. 2020), their detailed projections remain unclear. This is because retrograde tracers can be taken up by en passant fibers, leading to nonspecific labeling, and classical anterograde tracers lack cell-type specificity. To overcome these limitations, we used genetically modified mice and viral tracers to selectively label GABAergic neurons and examine their projections onto the MGB.

Injection of AAV2-hSyn1(S)-Flex-tdTomato-T2A-sypEGFP into each IC subregion of GAD67-Cre mice resulted in GFP-immunoreactive axon terminals primarily in the ipsilateral MGB (Figure 1; injection sites in Supplemental Information Figure S1), consistent with previous studies (Winer et al. 1996; Ito et al. 2009; Beebe et al. 2018; Anair et al. 2022), contralateral IC, and in the ipsilateral VNTB (Figure 3). Notably, GABAergic projections from the IC to the VNTB were observed for the first time in this study, although the general IC-to-VNTB projection has been previously described using a classical anterograde tracer (Faye-Lund 1986; Caicedo and Herbert 1993; Thompson and Thompson 1993; Vetter et al. 1993).

**Fig. 1.**
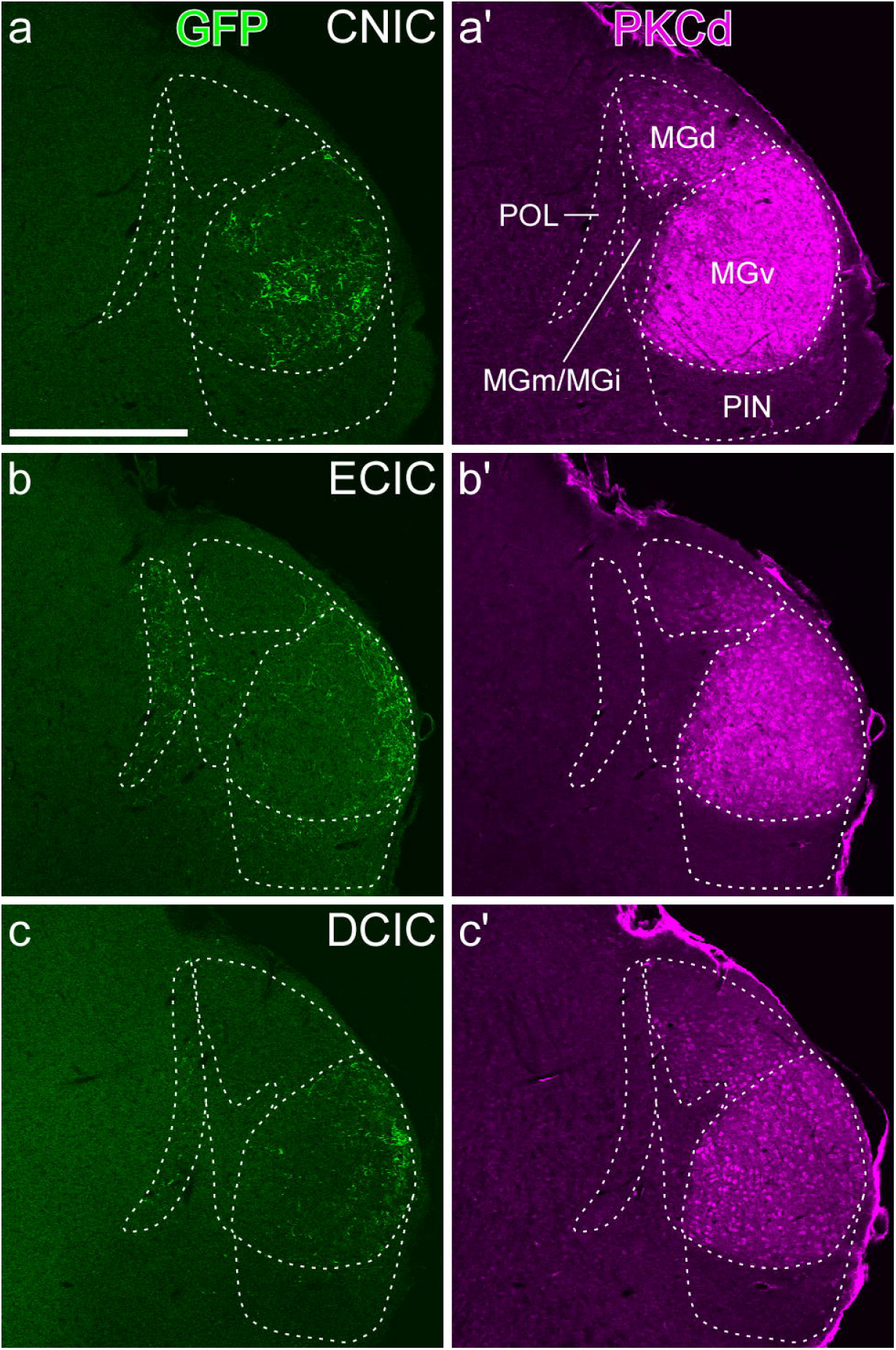
Axonal fibers of GABAergic neurons from IC subregions projecting to the MGB. AAV2-hSyn1(S)-Flex-tdTomato-T2A-sypEGFP was injected into each IC subregion of the GAD67-Cre mouse. a) Synaptophysin-GFP-positive fibers from GABAergic neurons in the CNIC were distributed mainly in the MGv. a’-c’) The boundaries of each MGB subdivision were delineated based on PKCδ immunoreactivity. b) Synaptophysin- GFP-positive fibers from GABAergic neurons in the ECIC were distributed mainly in the POL, MGv, PIN/PP. c) Synaptophysin-GFP-positive fibers from GABAergic neurons in the DCIC were distributed mainly in the MGv. The injection sites are displayed in the Supplementary Information Figure S1. Scale bar: 0.5 mm

We further analyzed the areas of GFP-immunoreactive axon terminals in the five subdivisions of the MGB according to their anatomical and neurochemical profiles (Llano and Sherman 2008; Márquez-Legorreta et al. 2016; Tomioka et al. 2023). The analyzed MGB subdivisions included the MGv, dorsal division of the MGB (MGd), medial (MGm) and internal (MGi) divisions of the MGB (Tomioka et al. 2023), posterior limiting nucleus (POL) (Ledoux et al., 1987; Cai et al. 2019; Liu et al. 2024a), posterior intralaminar nucleus, and peripeduncular nucleus (PIN/PP). PKCδ immunoreactivity clearly delineated these subregions, showing uniform labeling in MGv, a gradient pattern in MGd, and weak signals in the MGm/MGi and PIN/PP (Figure 1). Injections into the CNIC and DCIC resulted in GFP-immunoreactive axon terminals primarily in the MGv (Figures 1a,c and Figure 2), whereas ECIC injections labeled terminals primarily in the MGv, with weaker labeling observed in the PIN/PP and POL (Figure 1b and Figure 2). Statistical analysis using the Friedman test with Bonferroni correction demonstrated a significantly higher axonal density in the MGv than in the MGd and MGm/MGi following injection into the CNIC (p = 0.0365 and p = 0.0175, respectively) and DCIC (p = 0.0365 and p = 0.008, respectively) (Figure 2b). Considering that the MGv is a part of the lemniscal pathway, while the other MGB subregions belong to the non-lemniscal pathway, our findings suggest that GABAergic projection neurons from all IC subregions exert an inhibitory influence primarily on the lemniscal pathway, with those from the ECIC additionally targeting the non-lemniscal pathway. Notably, GABAergic neurons in the CNIC and ECIC also project to the ipsilateral VNTB and exert inhibitory effects (Figure 3).

**Fig. 2.**
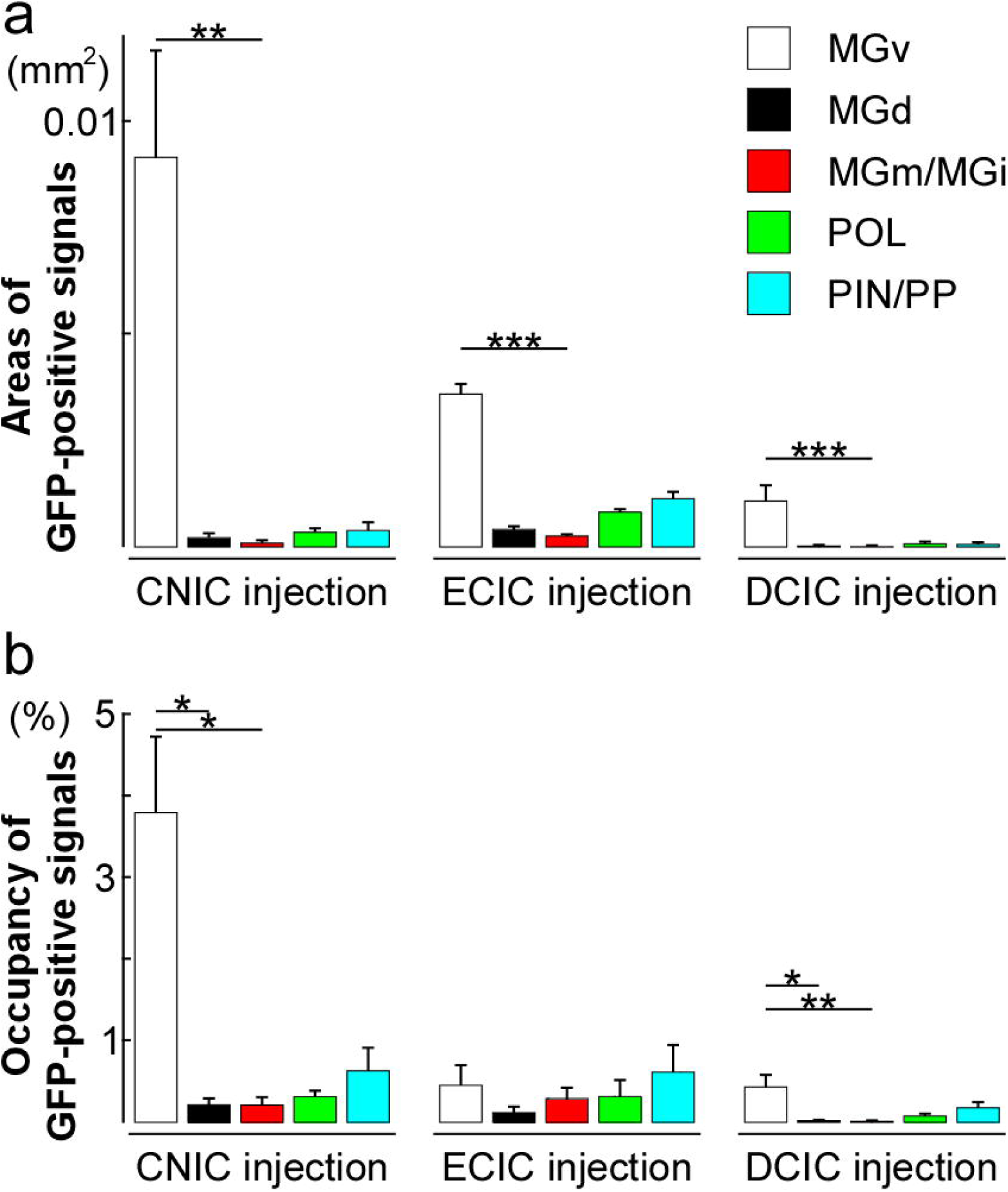
Areas and occupancy of GABAergic fibers from IC subregions projecting to the MGB. a) The area of axonal projections of CNIC GABAergic neurons within MGB was the largest in the MGv, with statistically significant differences observed among subregions (left panel; Friedman test; n=4). Similar GABAergic projections from the ECIC (middle panel) and DCIC (right panel) were observed. In multiple comparisons, a statistically significant difference was observed between MGv and MGm/MGi in axonal projections of GABAergic neurons from IC regions (Bonferroni correction). b) The occupancy of axonal projections of CNIC GABAergic neurons within MGB was also the largest in the MGv, with statistically significant differences observed among subregions (left panel; Friedman test; n=4). A similar pattern was observed in the GABAergic neurons of the DCIC (right panel), whereas no significant differences were observed in the axonal occupancy of ECIC GABAergic projections across MGB subregions. *p < 0.05; **p < 0.01; ***p < 0.005

**Fig. 3.**
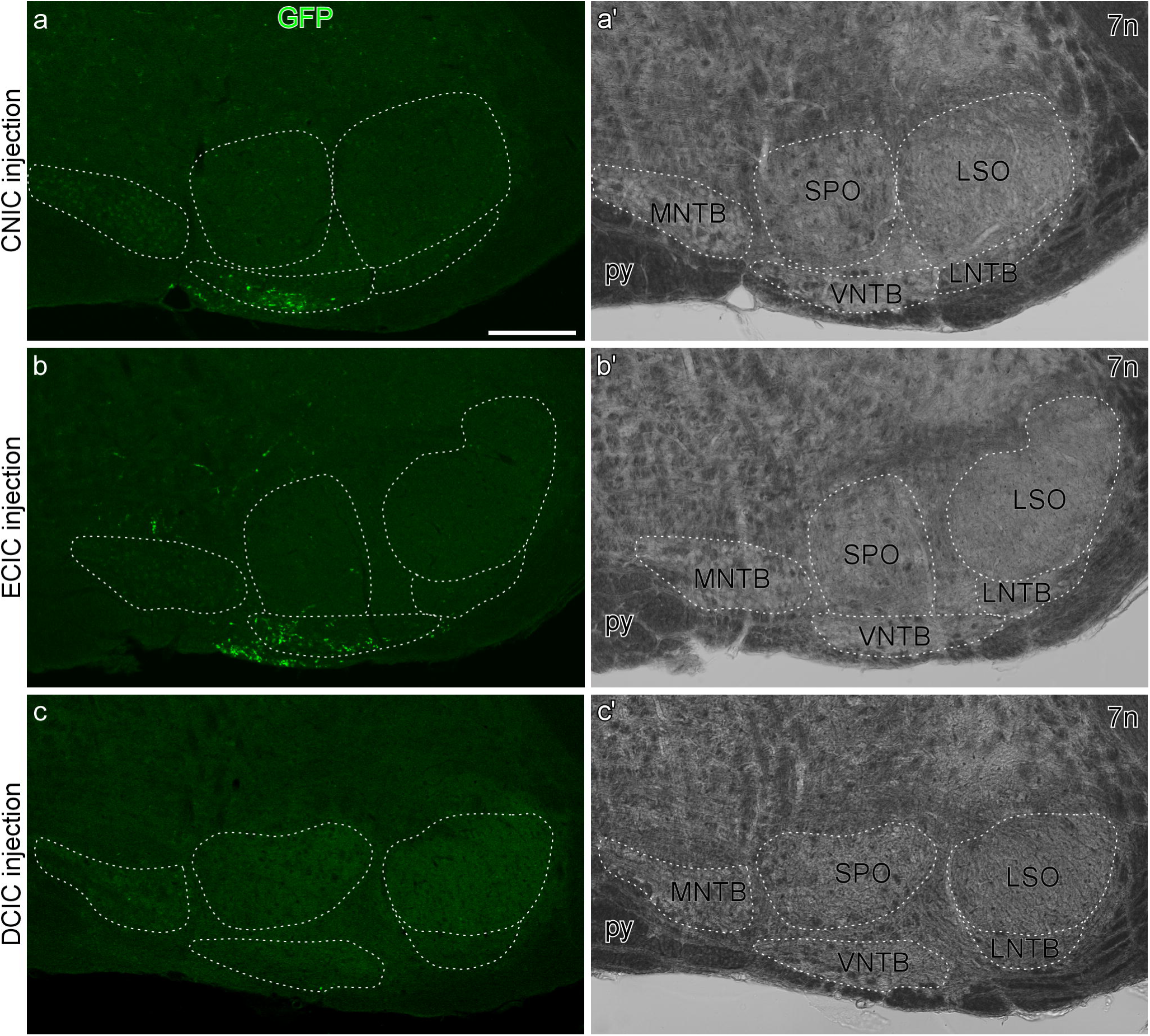
Axonal fibers of GABAergic neurons from IC subregions projecting to the SOC. a) Synaptophysin-GFP-positive fibers from GABAergic neurons in the CNIC were distributed mainly in the VNTB. b) Fibers from GABAergic neurons in the ECIC were mainly in the VNTB. c) Few fibers from GABAergic neurons in the DCIC were observed in the SOC. Scale bar: 200 µm

### Neurochemical profiles of GABAergic projection neurons in the IC

Accumulating evidence has demonstrated that GABAergic neurons in the IC are heterogeneous, based on their neurochemical profiles (Ouda and Syka 2012; Foster et al. 2014; Fujimoto et al. 2017; Schofield and Beebe 2019; Liu et al. 2024b). In the present study, we used GAD67-GFP mice to identify GABAergic neurons, thereby providing a more sensitive and reliable approach. Indeed, nearly all GFP-positive neurons in the IC of GAD67-GFP mice were GABA-immunoreactive (Ono et al. 2005). We examined the proportion of GABAergic neurons in the mouse IC (Figure 4), although previous studies estimated that they constitute approximately 20−30% of IC neurons in rats, cats, or bats (Oliver et al. 1994; Winer et al. 1995; Merchán et al. 2005; Fredrich et al. 2009). GFP-expressing neurons, used as markers for GABAergic neurons, were sparsely distributed in the IC, accounting for 10.2% of NeuN-immunoreactive neurons (5,249 NeuN-immunoreactive neurons analyzed from two animals): 12.0% in the CNIC, 11.4% in the ECIC, and 6.2% in the DCIC.

**Fig. 4.**
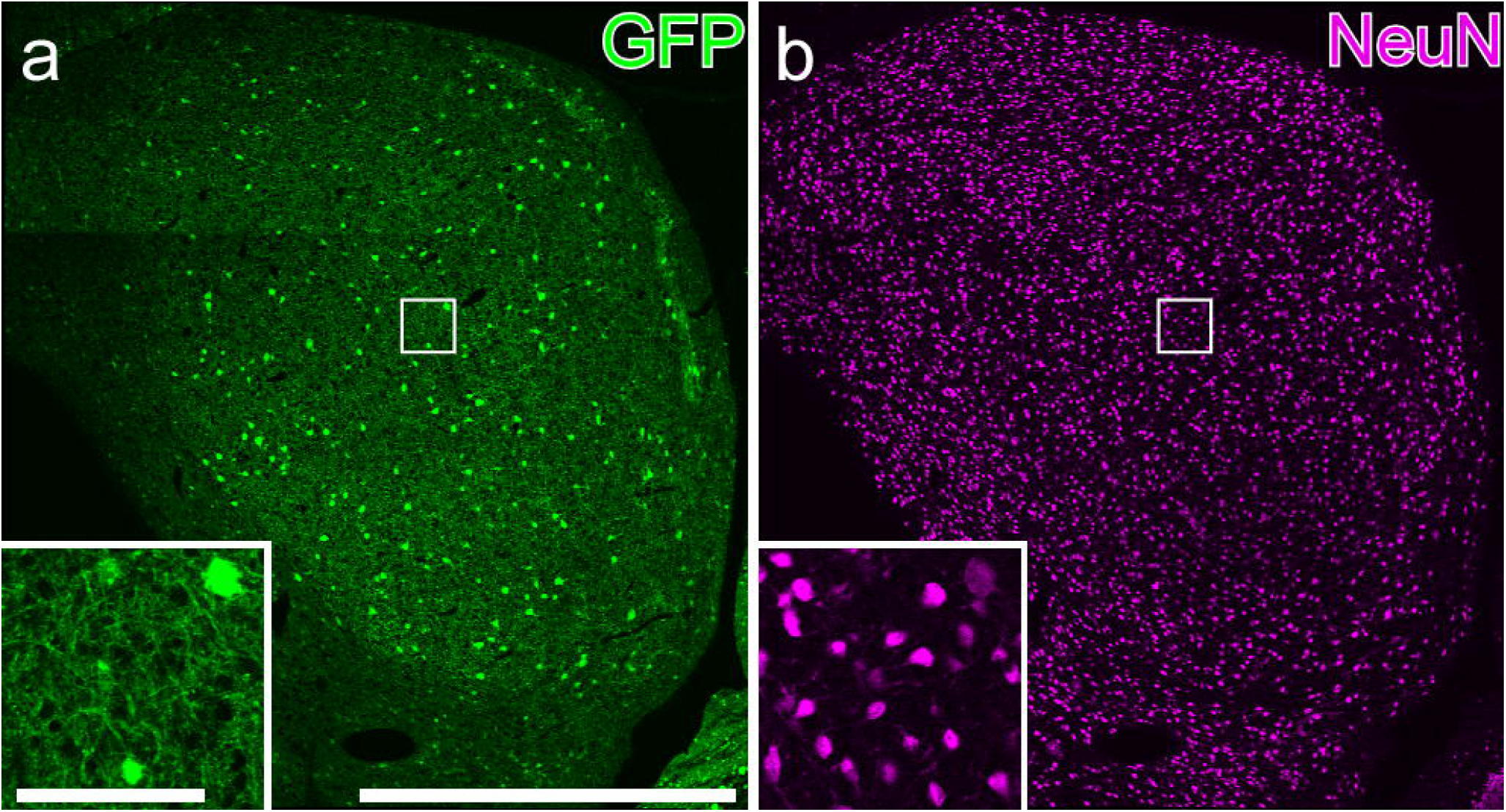
Proportion of GABA neurons in the IC. a) GFP fluorescence in GAD67-GFP mice exhibited the distribution of GABAergic neurons. b) NeuN immunoreactivity demonstrated the distribution of all neurons in the IC. The insets are the enlarged image from the rectangle area

Next, we examined the heterogeneity of GABAergic neurons in the IC using the neurochemical markers NPY, Kv3.1b, and PV in GAD67-GFP mice (Nakagawa et al. 1995; Weiser et al. 1995; Fredrich et al. 2009; Fujimoto et al. 2017; Silveira et al. 2020). NPY-immunoreactive neurons typically had relatively large somata and were mainly distributed in the ECIC (Figure 5a), accounting for 32.7% of the GFP-expressing GABAergic neurons (954 GFP-fluorescent neurons analyzed from three animals). Notably, 99% of the NPY-immunoreactive neurons were GFP-positive, indicating that NPY-positive neurons were GABAergic. Kv3.1b-immunoreactive neurons displayed variable soma sizes and were distributed throughout the IC (Figure 5b), accounting for 83.7% of the GFP-expressing GABAergic neurons (957 GFP-fluorescent neurons analyzed from three animals). Kv3.1b expression was observed in a substantial number of GFP-positive and GFP-negative neurons, suggesting that it is not specific to GABAergic neurons. PV-immunoreactive neurons exhibited variable soma sizes and were distributed throughout the IC (Figure 5c), accounting for 72.5% of GFP- expressing GABAergic neurons (960 GFP-fluorescent neurons analyzed from three animals). Notably, 96.5% of the PV-immunoreactive neurons were GFP-positive, indicating that PV-positive neurons were GABAergic. This finding is consistent with a previous report (Fredrich et al. 2009), but contrasts with more recent observations by Fujimoto et al. (2017) and Liu et al. (2024a) that reported PV-positive neurons in the IC include both GABAergic and glutamatergic subtypes.

**Fig. 5.**
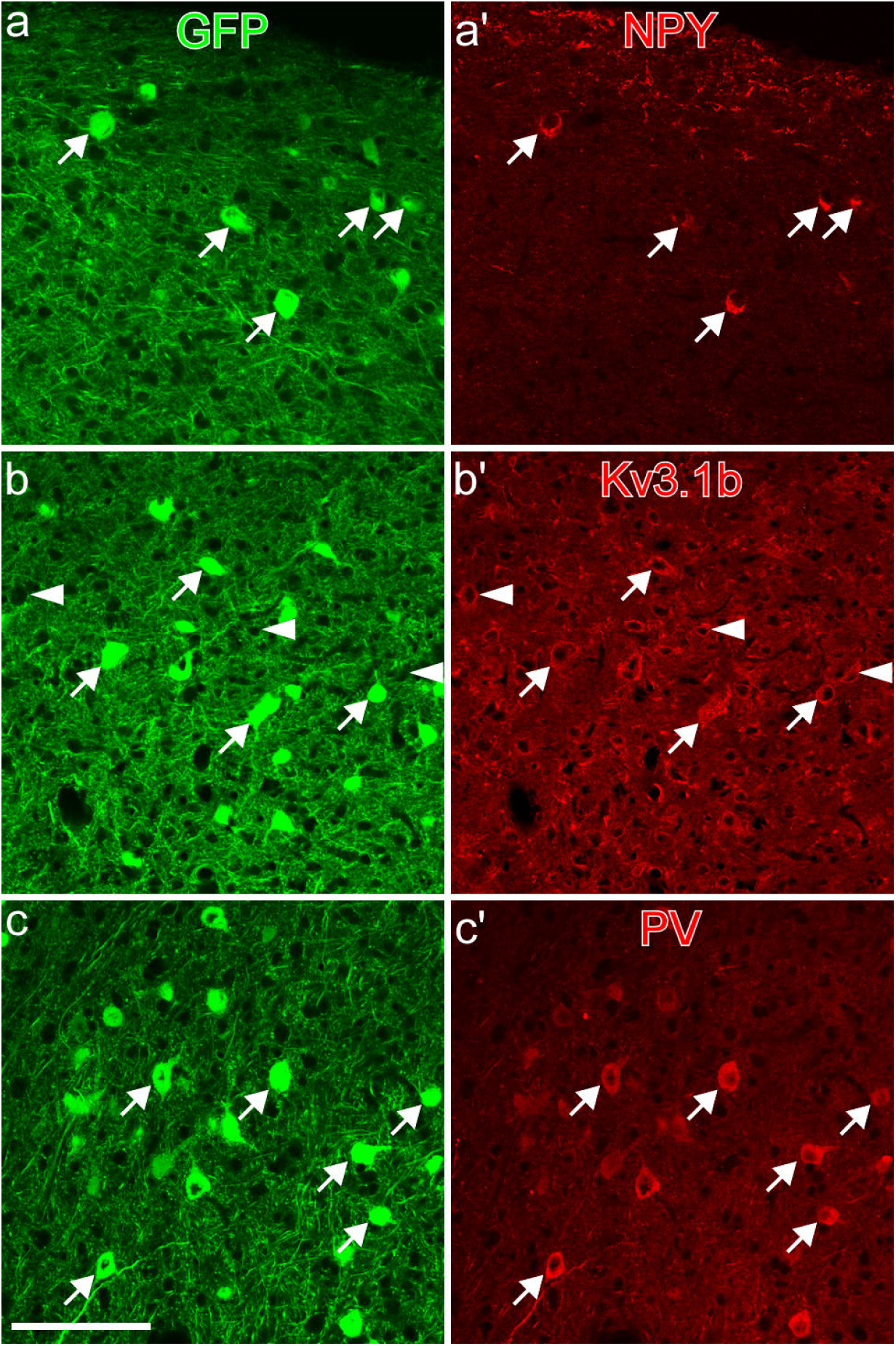
Neurochemical profiles of GABAergic neurons in the IC. GABAeric neurons express GFP in the GAD67-GFP mouse. a) Nearly all NPY-immunoreactive neurons were GABAergic (arrows). b) Most Kv3.1b-immunoreactive neurons were GABAergic (arrows), although some were non-GABAergic (arrowheads). c) Nearly all PV- immunoreactive neurons were GABAergic (arrows) and the vast majority of GFP- positive neurons were also PV-positive. Scale bar, 100 µm.

### Neurochemical profiles of GABAergic projection neurons in the IC targeting the MGB, contralateral IC, and SOC

GABAergic neurons with large somata were observed for all neurochemical markers (Figure 5). Because a large soma is characteristic of GABAergic projection neurons (Ito et al. 2009; Beebe et al. 2018), it is plausible that these markers are also expressed in GABAergic projection neurons. Based on our anterograde tracing experiments (Figures 1-3), we focused on three major projection targets: the MGB, IC, and SOC. Following FG injection into the MGB (Figure 6), FG- and GFP-double- labeled neurons were observed in both the ipsilateral and contralateral IC. A substantial proportion of retrogradely-labeled GABAergic neurons in the ipsilateral IC were immunoreactive for NPY, Kv3.1b, and PV: 58.8% for NPY (136 FG- and GFP-double– labeled neurons analyzed from four animals), 75.0% for Kv3.1b (98 FG- and GFP- double–labeled neurons analyzed from four animals), and 72.0% for PV (136 FG- and GFP-double–labeled neurons analyzed from four animals). Similarly, a high proportion of FG- and GFP-double-labeled neurons in the contralateral IC were positive for the following neurochemical markers: 84.2% for NPY (19 FG- and GFP-double–labeled neurons analyzed from four animals), 80.0% for Kv3.1b (15 FG- and GFP-double– labeled neurons analyzed from four animals), and 73.7% for PV (19 FG- and GFP- double–labeled neurons analyzed from four animals). To examine the commissural connectivity between ICs, FG was injected into the IC; retrogradely labeled FG- and GFP-double-–labeled neurons were subsequently observed in the contralateral IC. A majority of the retrogradely labeled GABAergic neurons in the contralateral IC were immunoreactive for the following neurochemical markers: 65.4% for NPY (78 FG- and GFP-double–labeled neurons analyzed from three animals), 75.0% for Kv3.1b (n = 98, three animals), and 92.3% for PV (78 FG- and GFP-double–labeled neurons analyzed from three animals). Following FG injections into the SOC, the vast majority of retrogradely labeled FG-positive neurons were observed in the ipsilateral IC and lacked GFP expression (232 FG-labeled neurons analyzed from three animals). In these experiments, all FG- and GFP-double-–labeled neurons exhibited NPY immunoreactivity (5 FG- and GFP-double–labeled neurons analyzed from three animals).

**Fig. 6.**
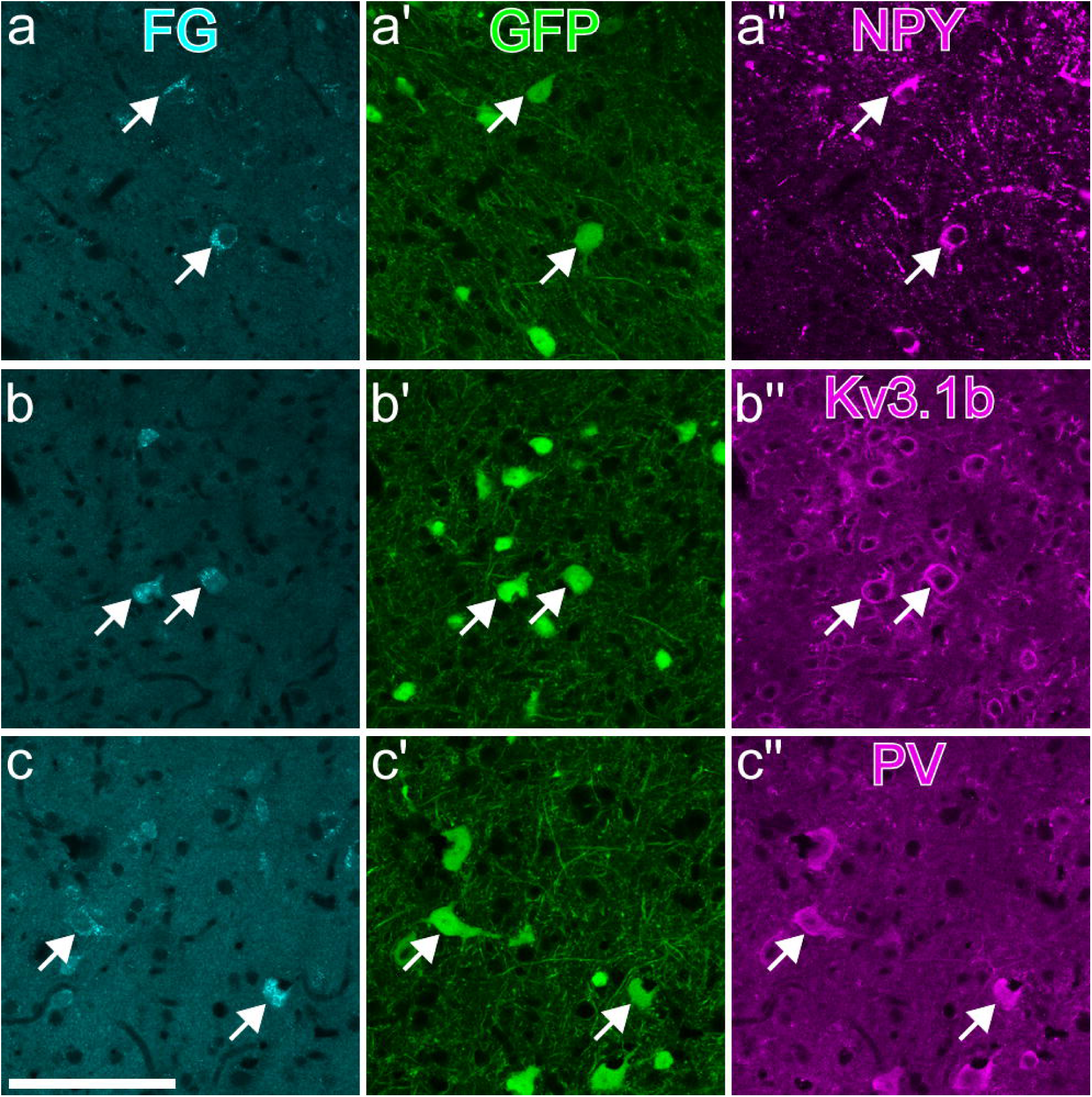
Neurochemical profiles of GABAergic projection neurons in the IC. Injection of the retrograde tracer FG into the MGB of GAD67-GFP mice resulted in FG-labeled neurons in the IC. a) FG-labeled GABAergic neurons were positive for NPY, Kv3.1b, and PV immunoreactivity (arrows). The injection sites are displayed in SI Figure S2. Scale bar: 100 µm

The proportion of GABAergic projection neurons at each FG injection site (MGB, IC, and SOC) was further investigated. In the MGB, IC, and SOC, 20%, 25%, and 2% of the FG-labeled neurons in the IC were identified as GABAergic, respectively. Given that GABAergic neurons constitute approximately 10% of the total neuronal population in the IC, the observation that they account for roughly 20% of IC–MGB projection neurons suggests that GABAergic neurons may contribute to this pathway more substantially than their overall prevalence would predict.

## Discussion

The present study builds on previous reports to provide a more detailed understanding of the anatomical and neurochemical profiles of GABAergic projection neurons in the mouse IC. By employing novel genetic applications, we characterized the distribution of axon terminals from GABAergic projection neurons in IC subregions. GABAergic neurons in the CNIC predominantly projected to the lemniscal region of the MGB, whereas GABAergic neurons in the ECIC targeted both the lemniscal and non- lemniscal regions, including the PIN/PP and POL. Unique GABAergic projections from the CNIC and ECIC to the VNTB were identified for the first time. Second, we demonstrated that GABAergic neurons comprised approximately 10% of all IC neurons, whereas approximately 20% of IC-MGB projection neurons were GABAergic. This difference suggests that GABAergic projection neurons play a particularly important role in the IC-MGB pathway. Third, we examined the neurochemical profiles of the GABAergic projection neurons in the IC. Notably, NPY and PV were reliable markers of GABAergic neurons, whereas Kv3.1b was expressed in both GABAergic and glutamatergic neurons.

Our findings posit the CNIC as a major source of GABAergic projection neurons in the mouse IC (Figure 7). These neurons would exert inhibitory effects primarily on the MGB and contralateral IC, and to a lesser extent on the ipsilateral VNTB. In addition to GABA, GABAergic projection neurons are likely to release the NPY neuropeptide, suggesting a modulatory role beyond classical inhibition (Silveira et al. 2020; Anair et al. 2022). Furthermore, the expression of Kv3.1b implies that these neurons may exhibit tonic firing properties, potentially providing sustained inhibitory inputs to target regions. The Kv3.1b channel protein enables expression of a fast-spiking phenotype by narrowing the spike width, shortening the refractory period, and ensuring complete membrane repolarization via a resurgent potassium current (Wang et al. 1998; Erisir et al. 1999; Rudy and McBain 2001; Labro et al. 2015; Tremblay et al. 2016). Taken together, these insights significantly advance our understanding of network dynamics within the central auditory system.

**Fig. 7.**
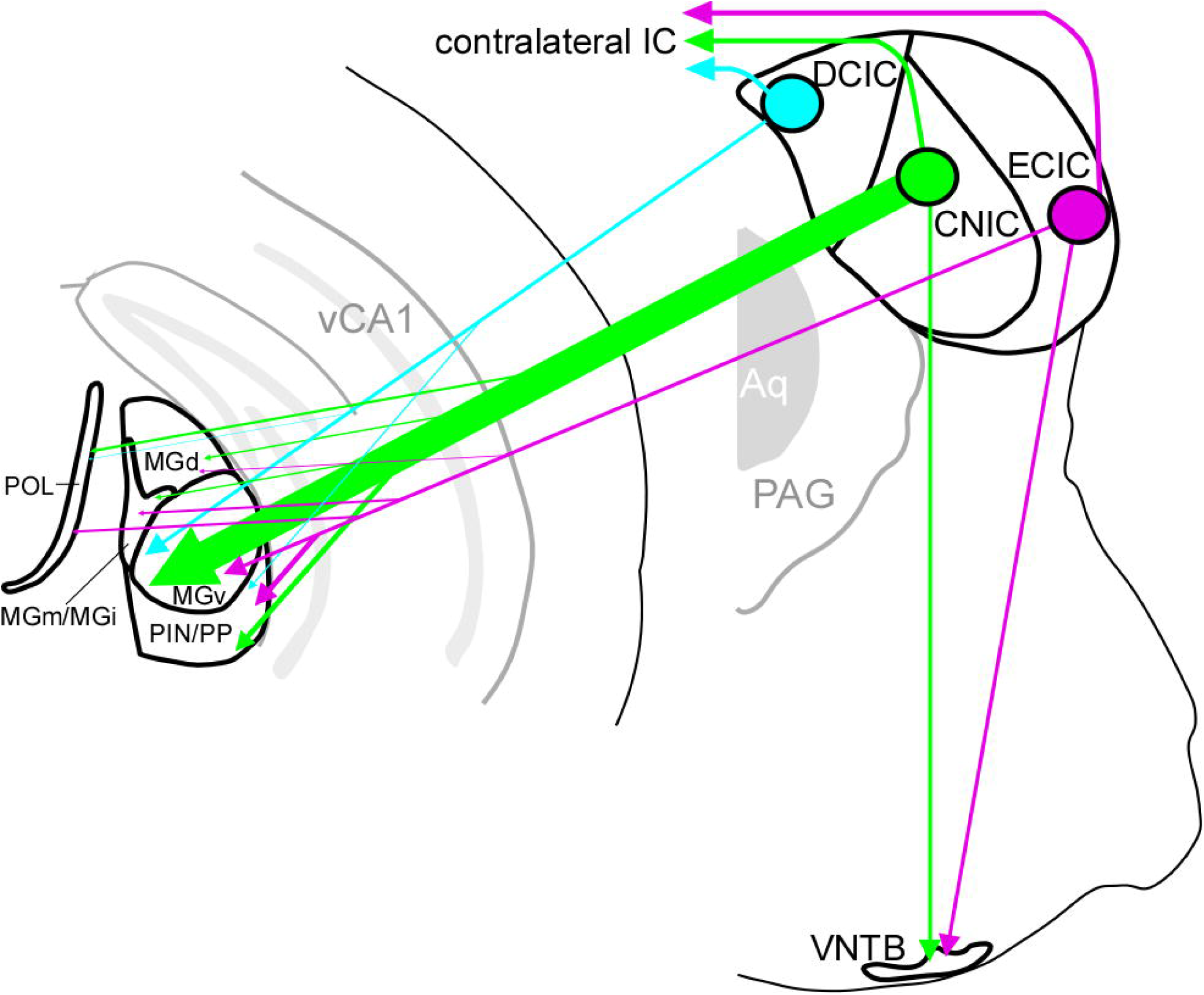
A schematic drawing of the GABAergic projection neurons in the IC. The thickness of the line corresponds to the occupancy of axon terminals. MGV received strong synaptic input from GABAergic projection neurons in all IC subregions, especially in CNIC. Note that, for the projection to the VNTB, the thickness of the line does not reflect the density of axon terminals

### Comparison with previous studies

Accumulating anatomical and neurochemical evidence indicates that even within a single brain nucleus structure, heterogeneous neuronal subgroups can be distinguished based on cellular or gene expression profiles, and that each neuronal subgroup often exhibits distinct projection patterns (Greig et al. 2013; Tasic et al. 2016; Jing et al. 2025). Moreover, it is also known that a single neuronal subgroup may project to multiple brain regions (Lin et al. 2018; Yuan et al. 2023; Tomioka et al. 2024). Therefore, we applied genetic techniques to selectively label GABAergic neurons and examined their axonal projections. Although GABAergic projection neurons in the IC appear to consist of several subtypes, as suggested by Beebe et al. (2018), our findings suggest the major subtypes are likely NPY-, Kv3.1b-, or PV-expressing neurons. To further enhance the detection of GABAergic neurons in GAD67-GFP mice, NeuN immunolabeling was concurrently employed. With NeuN immunolabeling, GABAergic neurons constituted approximately 10% of IC neurons, which is considerably lower than the roughly 20% reported in previous studies (Oliver et al. 1994; Merchán et al. 2005; Beebe et al. 2016). This discrepancy may be due to the improved detection of IC neurons by NeuN immunostaining. Furthermore, our combination of retrograde tracing and neurochemical marker analysis revealed that the same GABAergic neuronal subgroups project mainly to the MGB and minorly to the VNTB. These findings underscore the utility of genetically guided approaches for resolving cell type-specific connectivity.

Previous studies reported that PV-expressing neurons in the CNIC are GABAergic (Fredrich et al. 2009), whereas two recent studies reported that PV- expressing neurons in the IC include both glutamatergic and GABAergic subtypes (Fujimoto et al. 2017; Liu et al. 2024a). Our findings indicated that the vast majority of PV-positive neurons were GABAergic. This discrepancy may reflect differences in methodological approaches, such as the use of genetically modified mouse lines, immunohistochemical markers, or criteria for neuronal classification. Further studies using cell type-specific labeling combined with single-cell transcriptomics or electrophysiological profiling may help clarify the heterogeneity and functional roles of PV-expressing neurons in the IC.

### Functional implication of GABAergic projection neurons in the auditory system

Both glutamatergic and GABAergic neurons in the IC project to the MGB to form parallel excitatory and inhibitory pathways. Such dual-projection architecture is rarely observed in neural circuits other than the auditory system (Pollak et al. 2003; Tomioka et al. 2015; Weingarten et al. 2023; Maraslioglu-Sperber et al. 2024). This parallel organization may play a critical role in supporting the high temporal fidelity required for precise auditory processing. This structural characteristic may reflect the need of the auditory system to preserve and relay fine temporal information originating from lower brainstem nuclei such as the SOC, which encodes sound localization cues with sub-millisecond precision (Vonderschen and Wagner 2014). While the IC itself may not operate on the same microsecond scale, its role in integrating and transmitting such temporally precise inputs likely demands specialized circuit architecture.

To understand the unique contribution of GABAergic projection neurons in the IC, it is helpful to first consider the overall pattern of inputs to its primary target, the MGB. The MGB receives excitatory inputs from both the IC and layer 5/6 pyramidal neurons in the auditory cortex as well as inhibitory inputs from several regions, including the thalamic reticular nucleus (Kimura et al. 2007; Kimura and Imbe 2015), external segment of the globus pallidus (Tomioka et al. 2024), and IC. Notably, the MGB in mice contains very few intrinsic GABAergic neurons, indicating that inhibitory control largely depends on extrinsic sources. Inputs from the IC provide ascending (feed-forward) pathways, whereas those from the auditory cortex, TRN, and GPe represent descending (feedback) pathways. This feed-forward input is associated with real-time auditory information processing, whereas higher-order auditory areas that exhibit broad-frequency tuning and slower response latencies are thought to play a role in integrating auditory information over longer timescales (Joris et al. 2004). Thus, GABAergic neurons in the IC are directly involved in the transmission of auditory information. One proposed mechanism is that GABAergic projection neurons deliver feed-forward inhibitory postsynaptic potentials (IPSPs) to the MGB before excitatory postsynaptic potentials (EPSPs) arrive from simultaneously firing glutamatergic IC neurons, thereby sharpening the temporal precision of MGB responses (Ito et al. 2009). Another potential function, suggested by studies in other brain regions, is the formation of a temporal window in which inhibitory inputs regulate the activity of postsynaptic neurons (Ouda and Syka 2012; Tremblay et al. 2016). This mechanism is believed to refine the timing of neuronal firing by defining the precise windows during which postsynaptic neurons are permitted to fire. Since the latency of EPSPs in response to acoustic stimuli is typically shorter than that of IPSPs in MGB neurons (Yu et al. 2004), the latter mechanism, temporal window formation, may be more likely. Notably, the expression of Kv3.1b and PV in GABAergic projection neurons resembles cortical fast- spiking GABAergic neurons, which are known to enhance temporal precision by establishing time windows for postsynaptic firing (Tremblay et al. 2016). This similarity implies that GABAergic projection neurons in the IC may have a comparable function in refining auditory timing. A comprehensive understanding of the function of these GABAergic projection neurons in auditory processing requires further anatomical and physiological investigations. Elucidating how these neurons interact with their glutamatergic counterparts within this parallel architecture could reveal the fundamental principles of the auditory network organization unique to this sensory modality.

In addition to their projections to the MGB, this study revealed that the GABAergic projection neurons in the IC send descending projections to the ipsilateral VNTB. Although the presence of GABAergic terminals in the VNTB suggests the potential for inhibitory modulation, our results indicate that this projection is relatively minor. It is likely that glutamatergic neurons contribute significantly to the IC–VNTB connection, providing one of the excitatory inputs to this region alongside major inputs from the cochlear nucleus. Given that VNTB neurons are predominantly inhibitory, either glycinergic or GABAergic, and project mainly to the contralateral dorsal cochlear nucleus (DCN) and ipsilateral IC(Shore et al. 1991; Vetter et al. 1993; Warr and Beck 1996; Ostapoff et al. 1997; Ito et al. 2011; Gómez-Martínez et al. 2025), IC–VNTB projections may contribute to shaping auditory processing at multiple levels of the auditory system. Importantly, these VNTB projections are topographically organized to process auditory information from the same side of space.

Thus, GABAergic projection neurons in the IC could exert a widespread influence on auditory function through direct inhibitory inputs to the MGB, contralateral IC, and VNTB via glutamatergic projection neurons. Such coordinated activity between inhibitory and excitatory pathways may contribute to the precise temporal and spatial shaping of auditory information across multiple levels of the auditory system.

## Acknowledgments

We thank the members of W.J.S’s laboratory for the discussion. We also thank Ryoko Mine at the Center for Metabolic Regulation of Healthy Aging, Faculty of Life Sciences, Kumamoto University, and the members of the Center for Animal Resources and Development, Kumamoto University for their technical support. This work was supported by JSPS KAKENHI Grant Number 24K09697 (R.T.), 22K06433, (W.J.S.), by Kawai Foundation for Sound Technology & Music (R.T.), and by Zoom Group Foundation for the Promotion of Science Grant (R.T.).

## Abbreviations

Aq: Aqueduct
CN: cochlear nucleus
CNIC: central nucleus of inferior colliculus
DCIC: dorsal cortex of inferior colliculus
ECIC: external cortex of inferior colliculus
IC: inferior colliculus
LNTB: lateral nucleus of trapezoid body
LSO: lateral superior olive
MBG: medial geniculate body
MGd: dorsal division of MGB
MGi: internal division of MGB
MNTB: medial nucleus of trapezoid body
MGm: medial division of MGB
MSO: medial superior olive
PAG: periaqueductal gray
PIN: posterior intralaminar nucleus
POL: posterior limiting nucleus
PP: peripeduncular nucleus
SOC: superior olivary complex
SPO: superior paraolivary nucleus
vCA1: ventral field CA1 of hippocampus
VNTB: ventral nucleus of trapezoid body

## Statements and Declarations

### Funding

This work was supported by JSPS KAKENHI Grant Number 24K09697 (R.T.), 22K06433, (W.J.S.), by Kawai Foundation for Sound Technology & Music (R.T.), and by Zoom Group Foundation for the Promotion of Science Grant (R.T.).

### Competing Interests

The authors declare no competing financial interest.

### Author contributions

Conceptualization: Ryohei Tomioka; Formal analysis: Ryohei Tomioka; Funding acquisition: Ryohei Tomioka, Wen-Jie Song; Investigation: Ryohei Tomioka; Methodology: Kenta Kobayashi; Resources: Wen-Jie Song; Visualization: Ryohei Tomioka; Writing – original draft: Ryohei Tomioka; Writing – review & editing: Ryohei Tomioka, Kenta Kobayashi, Wen-Jie Song.

### Data availability

Data that support the findings of this study are available from the corresponding authors upon reasonable request.

### Ethics approval

All protocols were approved by the Kumamoto University Animal Experimentation Committee (reference number: A2023-014R1)

**SI Fig. S1** Representative AAV injection sites. AAV2-hSyn1(S)-Flex-tdTomato-T2A- sypEGFP expresses tdTomato in a Cre-dependent manner. Injection sites illustrated for the a) CNIC, b) ECIC, and c) DCIC. Scale bar: 0.5 mm

**SI Fig. S2** Representative FG injection sites. The injection sites are the a) MGB, b) CNIC, and c) SOC. Scale bar: 0.5 mm

## Notes

### Competing Interest Statement

The authors have declared no competing interest.

