## Supplementary figures and images for "Anatomical and Neurochemical Profiles of GABAergic Projection Neurons in the Mouse Inferior Colliculus"

### Supplemental Figure 1

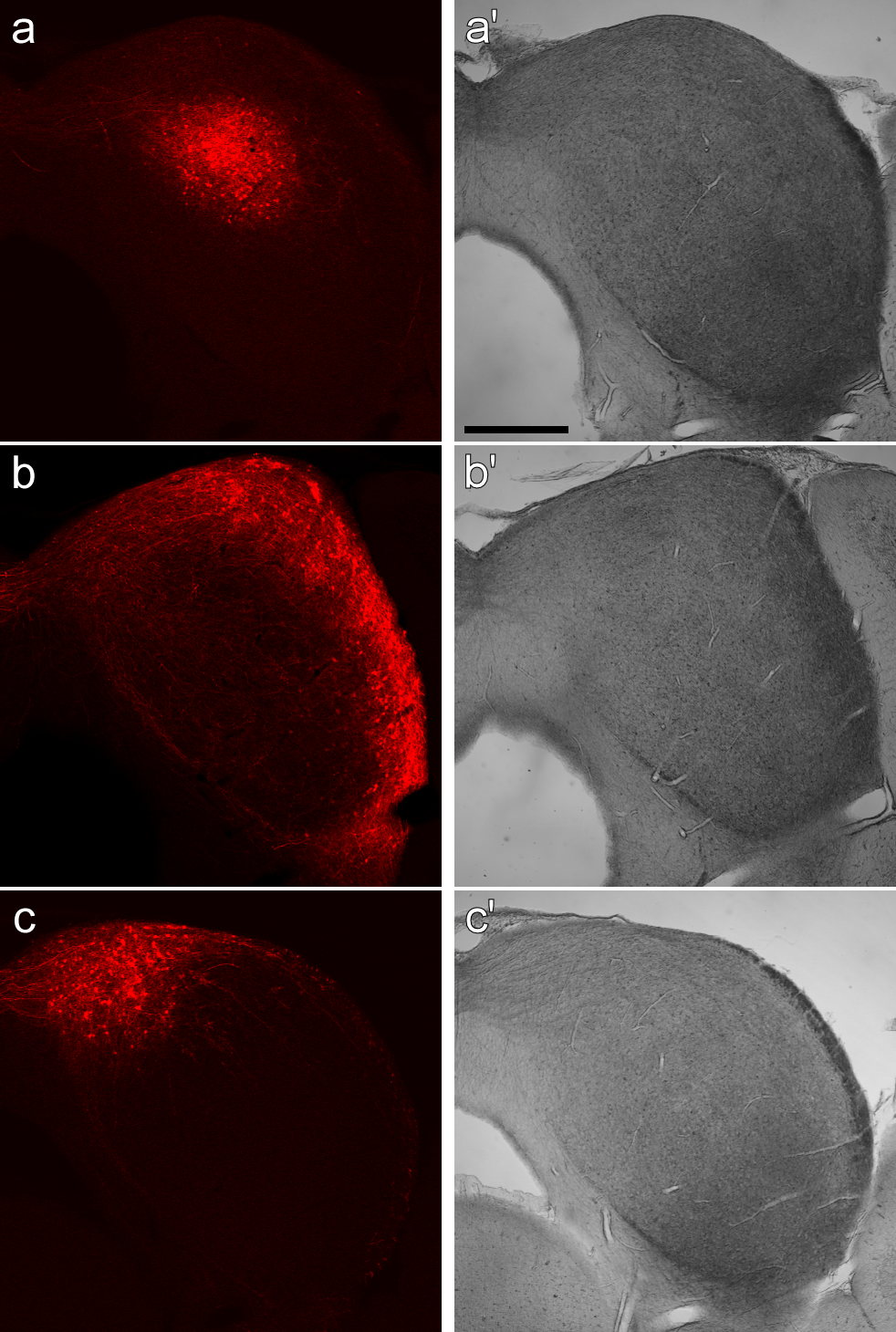

### Supplemental Figure 2

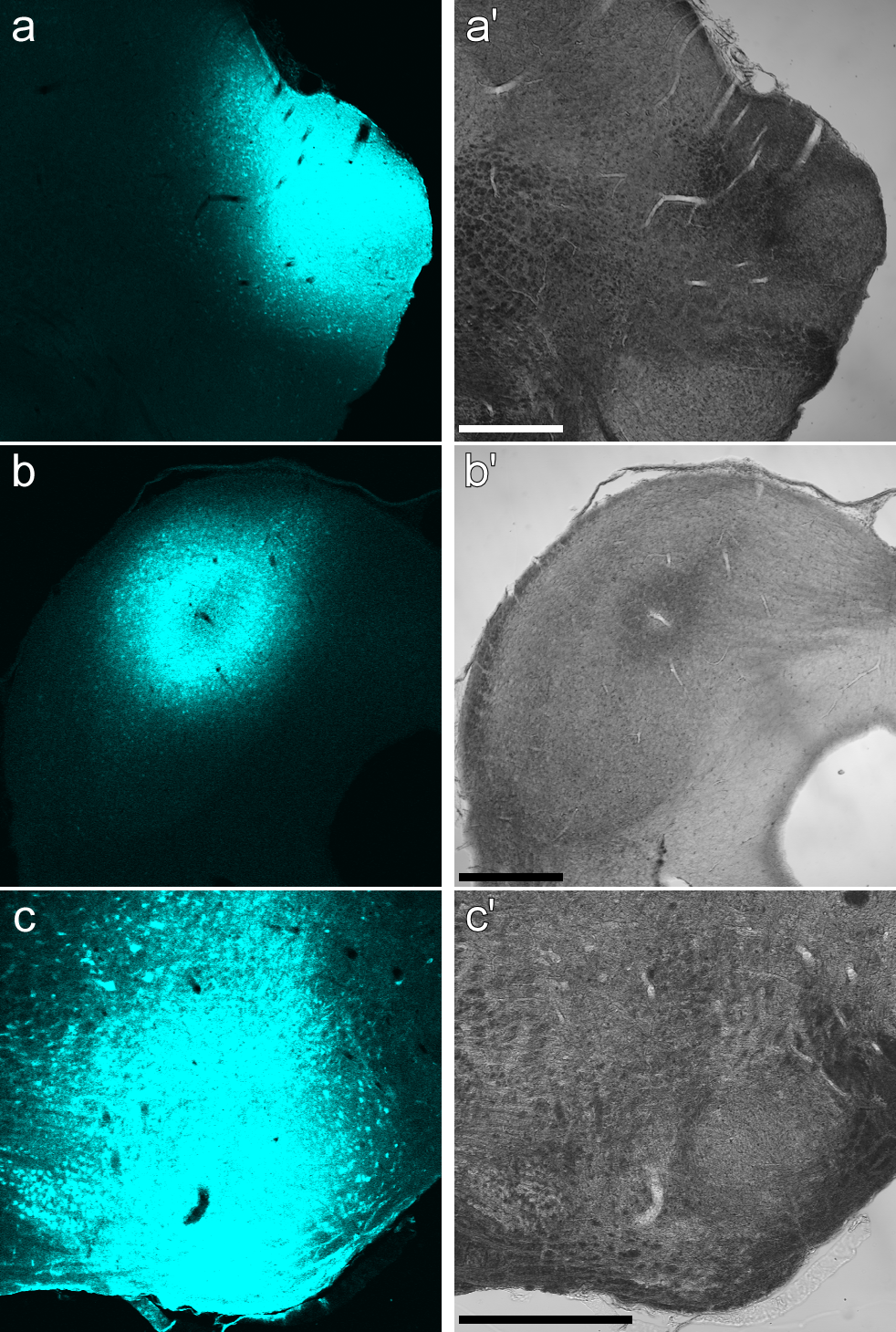
